# Self-healing dyes for super-resolution microscopy

**DOI:** 10.1101/373852

**Authors:** Jasper H. M. van der Velde, Jochem Smit, Michiel Punter, Thorben Cordes

## Abstract

In recent years optical microscopy techniques have emerged that allow optical imaging at unprecented resolution beyond the diffraction limit. Up to date, photostabilizing buffers are the method of choice to realize either photoswitching and/or to enhance the signal brightness and stability of the employed fluorescent probes. This strategy has, however, restricted applicability and is not suitable for live cell imaging. In this paper, we tested the performance of self-healing organic fluorophores with intramolecular photostabilization in super-resolution microscopy with targeted (STED) and stochastic readout (STORM). The overall goal of the study was to improve the spatial and temporal resolution of both techniques without the need for mixtures of photostabilizing agents in the imaging buffer. Due to its past superior performance we identified ATTO647N-photostabilizer conjugates as suitable candidates for STED microscopy. We characterize the photostability and resulting performance of NPA-ATTO647N oligonucleotide conjugates in STED microscopy. We find that the superior photophysical performance results in optimal STED imaging and demonstrate the possibility to obtain single-molecule fluorescent transients of individual fluorophores while illuminating with both the excitation- and STED-laser. Secondly, we show an analysis of photoswitching kinetics of self-healing Cy5 dyes (comprising TX, COT and NPA stabilizers) in the presence of TCEP- and cysteamine, which are typically used in STORM microscopy. In line with previous work, we find that intramolecular photostabilization strongly influences photoswitching kinetics and requires careful attention when designing STORM-experiments. In summary, this contribution explores the possibilities and limitations of self-healing dyes in super-resolution microscopy of differing modalities.

## 1. Introduction

In recent years, pioneering developments in far-field fluorescence microscopy have revolutionized optical imaging. Commonly known as “super-resolution microscopy” (or “nanoscopy”), they allow for visualization of biological structures beyond the physical diffraction-limited resolution of ~250 nm^1–8^. These super-resolution microscopy techniques all rely (to some extent) on ON/OFF-switching of fluorescence emission, *e.g.*, photoswitching. The available techniques can be grouped into localization-based approaches using video-type fluorescence microscopy such as STORM^1–3^, PALM^4^ and PAINT^5^ or targeted readout that is often realized in confocal setups, e.g., STED^6^, GSD^7^ and RESOLFT^8^.

As for any imaging approach that uses fluorescence as ‘molecular contrast’ agent, the performance and properties of the labels are of utmost importance. It became clear that one of the bottlenecks for high spatial resolution is the physical size of the label and its bio-linker^9,10^. Furthermore, signal- and photostability of the employed fluorophores play a role next to functional aspects such as photoswitching capabilities^1–8^. Whereas signal- and photostability determine the image contrast and allows for dynamic and time-lapse modalities, photoswitching is required to overcome the diffraction-limited resolution^11–14^. Up to date, often a mixture of buffer additives is used to improve the photophysical properties (triplet-state quenchers, antioxidants and others) and at the same time enabling photoswitching (thiol derivatives)^15,16^. During the past years, it became possible to use virtually any kind of synthetic organic fluorophore, fluorescent protein and even some semi-conductor nanocrystals for super-resolution microscopy.

More recently, intramolecular triplet-state quenching^17–19^ of synthetic organic dyes (“self-healing”^20–22^) emerged as an alternative strategy to achieve high photostability^20,23,24^. This approach overcomes important caveats of adding photostabilizers to the imaging buffer^20, 23–29^. For example, these conditions are often incompatible with biological requirements of living cells. Previous studies by our group demonstrated that self-healing dyes allow for STED-type imaging with a higher number of accessible successive images due to an increased photostability in fixed mammalian cells^24^, while the Blanchard lab showed cellular imaging with self-healing dyes for various standard fluorescence microscopy techniques.^20,25^

While the improvement of fluorophore signal and count-rate are directly linked to photostability, the design of intrinsically blinking fluorophores (ON/OFF switching) remains a challenge and has only been realized successfully for a handful of examples^30,31^. In this contribution, we explored the achievable spatial and temporal resolution of STED- and STORM-type microscopy techniques using self-healing dyes based on ATTO647N and Cy5. For this, we studied photostability, spatial resolution and brightness of single ATTO647N molecules in confocal and STED microscopy. We show that a single photostabilizer-ATTO647N derivative can provide MHz photon count rates, making the dye an ideal candidate for STED microscopy. Single molecule STED measurements of ATTO647N and NPA-ATTO647N revealed a significantly improved photostability of NPA-ATTO647N under STED imaging conditions. This allowed for multiple successive STED images of the same imaging area, even at elevated intensities of the STED laser. With NPA-ATTO647N a reduction of the point-spread-function down to 35 nm could be observed – values that are similar to resolution achieved with solution-based additives based e.g., on ROXS^10^. Additionally, for ATTO647N-photostabilizer derivatives it was possible to obtain fluorescent time traces of single dyes while illuminating with both the confocal excitation- and STED-laser. Finally, we explored the effects of intramolecular stabilizers (nitrophenylalanine (NPA), trolox (TX), cyclooctatetraene (COT)) on photoswitching kinetics of Cy5 for use in STORM-type microscopy. As a photoswitching agent we used reductive caging of fluorophores with tris(2-carboxyethyl)phosphine (TCEP) and the standardized condition in STORM, which is cysteamine-based blinking of cyanine dyes. We find a strong influence of the stabilizers on the photoswitching kinetics of Cy5 that have to be taken into account whenever self-healing dyes are used in STORM-type imaging.

## 2. Results

### Photostabilizer-dye conjugates for increased photostability in STED microscopy

A key element for optical super-resolution is the control of fluorescence emission signal (stable vs. blinking) as well as photostability. While STED microscopy requires stable and long lasting emission, photoswitching^6,7,32^ is essential to PALM/STORM^1–4^. Here, we benchmarked the performance of ATTO647N, a widely-used carbopyronine fluorophore, on double-stranded DNA against its photostabilizer-dye conjugate in STED microscopy. First, we investigated the photophysical behaviour of ATTO647N using single-molecule fluorescence microscopy. In the absence of oxygen ATTO647N shows photobleaching times on the order of seconds to minutes accompanied by frequent on/off blinking on the millisecond timescale (Figure 1a). This blinking can be assigned to a triplet-related dark-state (off-state lifetime is 29 ± 5 ms)^33^. When bound to the photostabilizer NPA the photophysical behaviour of ATTO647N changes and bright non-blinking molecules can be observed (Figure 1b). This can also be seen in the autocorrelation curve where the triplet-related amplitude is strongly reduced (Figure 1a and b). As reported previously^33^ the total number of detectable photons increases substantially. These features render NPA-ATT647N a suitable dye for STED microscopy, where photostability, brightness and signal to noise ratio (SNR) are important photophysical parameters.

**Figure 1:**
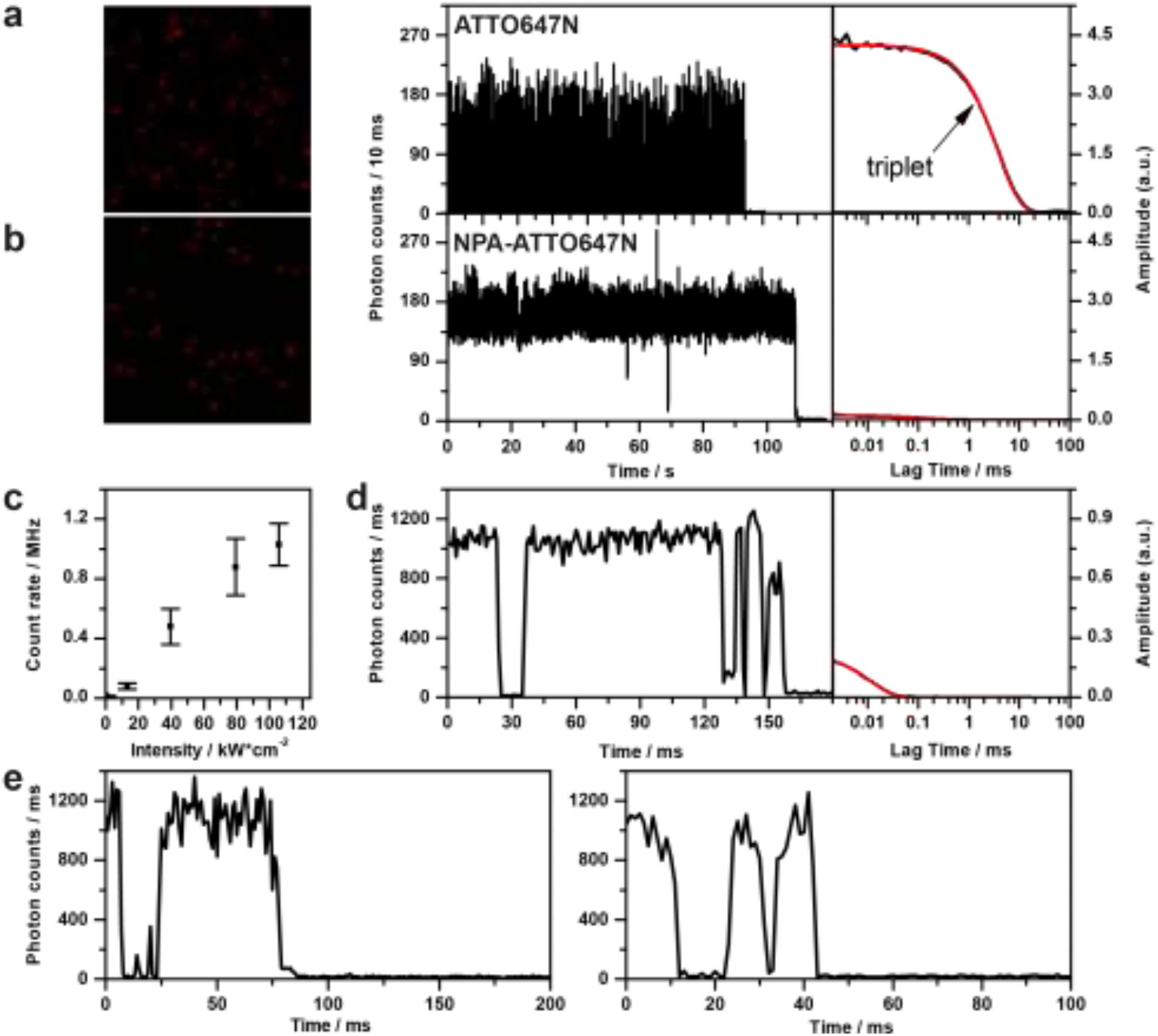
Photophysical characterization of NPA-ATTO647N with confocal microscopy. (**a**, **b**) A representative confocal overview image (10 x 10 μm, 50 nm pixel size, 2 ms per pixel) with spots from individual immobilized fluorophores (left), a fluorescence time trace (middle) and the corresponding autocorrelation decay (black) and fits (red) for ATTO647N (**a**) and NPA–ATTO647N (**b**). Image intensity scale from 5 to 100 counts, excitation intensity of ~0.66 kW/cm^2^ at 640nm. (**c**) Intensity dependence of the fluorescent count rate of NPA-ATTO647N. (**d**) A confocal trace of NPA-ATTO647N at 100 kW/cm2 (excitation at 640 nm) with corresponding autocorrelation function (black) and fit (red). (**e**) Additional confocal traces of NPA-ATTO647N at 100 kW/cm^2^ (excitation at 640 nm).

In order to quantify the maximum brightness and obtainable signal-to-background (SNB) and signal-to-noise (SNR) ratio, we increased the excitation intensity from moderate levels (50 W/cm^2^) up to 50-100 kW/cm^2^ (Figure 1c). Under these conditions, a single NPA-ATTO647N fluorophore emitted with MHz count rates (Figure 1c and d). To the best of our knowledge, such high values have previously only been obtained with the help of plasmonic effects^34,35^ or for significantly shorter observation times^36^ than is demonstrated here (>50-100 ms). Count rates of 1000 counts/ms allow for binning of the fluorescence signal with 50 μs resulting in SNR values of ~4-5. At intensities of 100 kW/cm^2^, an amplitude is found in the autocorrelation function with an off-state lifetime of < 10 μs (Figure 2d). It is likely associated with an increasing population of the triplet-state and might reduce photon output and SNR at higher excitation intensities.

**Figure 2.**
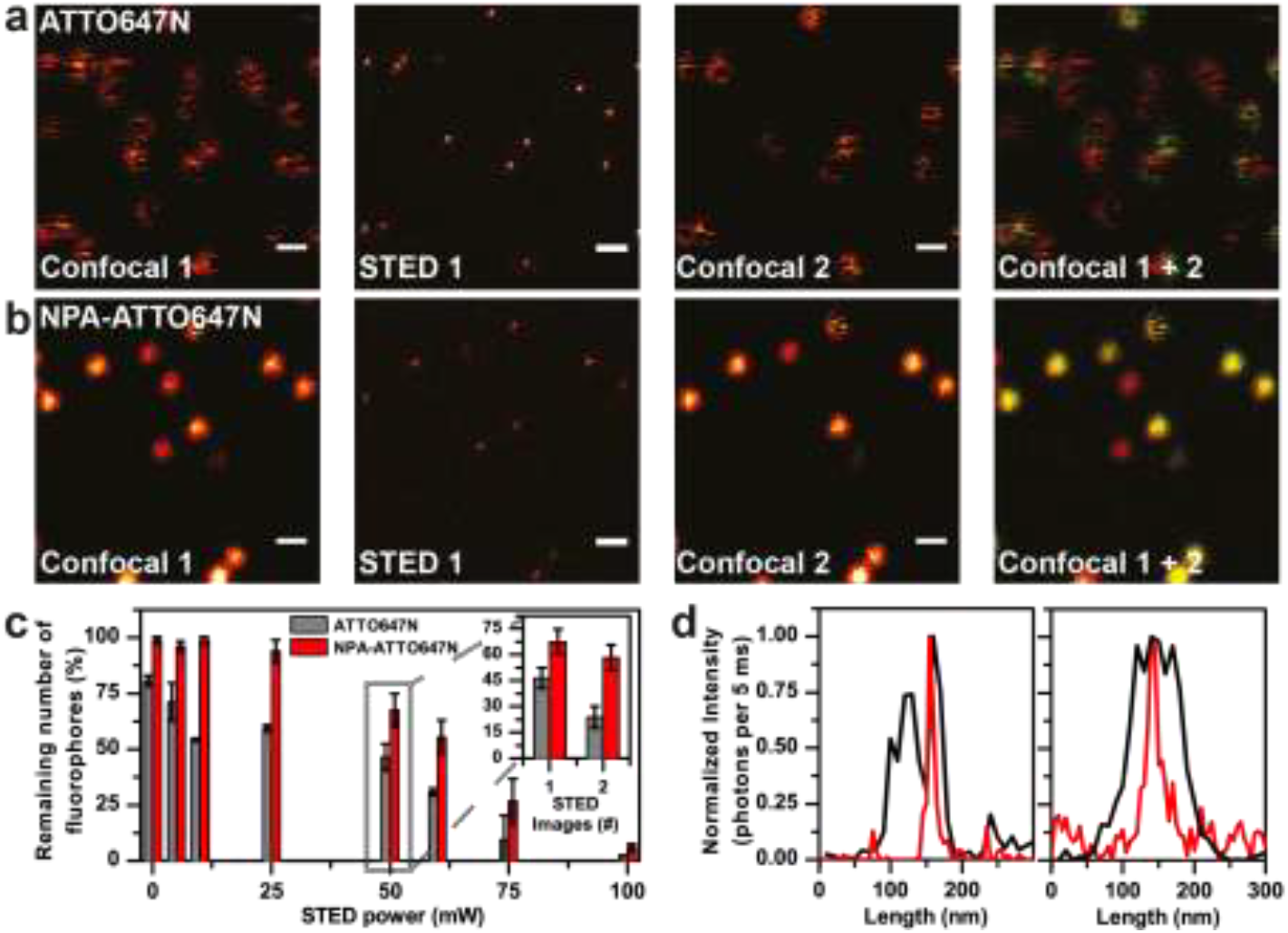
STED microscopy of individual ATTO647N derivatives. Alternating confocal, STED and confocal images of single (**a**) ATTO647N and (**b**) NPA-ATTO647N labelled oligonucleotides. Confocal 1+2 is an overlaid image of the first and second confocal image to visualize the number of remaining single molecules after a single STED image acquisition. (**c**) Quantitative analysis of the remaining number of fluorophores in a confocal image after acquiring a STED image at different STED laser powers for ATTO647N and NPA-ATTO647N. At 50 mW STED laser power the remaining number of fluorophores in a confocal image was analysed after acquiring one or two STED images (insert). (**d**) Line profiles of a single ATTO67N (left) and NPA-ATTO647N (right) molecule in confocal (black) and corresponding STED (red) image. The confocal images were recorded at 640 nm excitation and 3.4 kW cm^-2^. The STED images were recorded with 640 nm excitation at 3.4 kWcm^-2^ complemented with a STED beam at 765 nm with 50 mW at the sample (panel a and b). Scale bars are set to 500 nm. All data was recorded with a sample in aqueous PBS buffer at pH 7.4 in the absence of oxygen.

Next, STED microscopy was used to test whether NPA-ATTO647N also enhances the photostability under STED imaging conditions. Figure 2 shows alternating confocal and STED images of ATTO647N and NPA-ATTO647N under deoxygenated conditions. In the sequence of images, at first a confocal image was taken. Subsequently, an image of the same area was recorded with the STED beam overlaid to the confocal excitation beam. Finally, a second confocal image was recorded and compared to the first confocal image. In Figure 2a and b the analysis method was adapted from Kasper *et al.*^10^ From the composite image (confocal 1 + 2), it can be seen that a significant population of the ATTO647N molecules were photobleached after one scan of the field of view with the STED laser. On the contrary, the majority of NPA-ATTO647N molecules survived even a second cycle of confocal/STED imaging (Figure 2b and Figure 2c, inset), which is seen by the number of yellow spots in the confocal 1 + 2 image (Figure 2a and b). From this can be concluded that intramolecular photostabilization remains effective not only for labels in fixed cells^24^ but also for single-molecules under STED imaging conditions and high excitation powers.

For a quantitative analysis, the same series of alternating confocal and STED images was repeated at different powers of the STED beam (Figure 2c). At low powers it can be seen that already ~25-45% of individual ATTO647N fluorophores have photobleached after or during the acquisition of one STED image, whereas for NPA-ATTO647N almost all molecules are still fluorescent after STED imaging. Upon increasing the laser power of the STED beam, i.e., up to 60 mW, still >60% of the NPA-ATTO647N molecules do not photobleach after one STED-image. Further increasing the STED laser power photobleaches almost all (>90%) of the ATTO647N fluorophores, which is in contrast to NPA-ATTO647N where >25% of the molecules are still fluorescent. At 100 mW of STED power still ~6% of the single NPA-ATTO647N fluorophores survive the STED image acquisition. In contrast a minor fraction of only <2% of the ATTO647N fluorophores can still be detected. We assume here that in general a higher power of the STED laser results in an increased resolution^6,10,32^.

In a next step, moderate STED laser intensities of 50 mW were used to acquire multiple subsequent confocal and STED images. In Figure 2c (inset) the remaining number of fluorophores after one and two confocal-STED-confocal imaging cycles is shown. Again the difference between NPA-ATTO647N and its parent fluorophore is substantial with ~60% versus ~20% survival, respectively. This imaging mode with photostabilizer-dye conjugates allows for measuring multiple successive STED images of the same area with a drastically reduced loss of fluorescent signal (increased photostability), potentially enabling time-lapse STED imaging.

Next, the resolution dependence was studied as a function of the intensity of the STED laser. In the confocal image a Gaussian of the microscope’s point-spread function resulted in a FWHM of 225 ± 26 nm for NPA-ATTO647N (Figure 3a).

**Figure 3.**
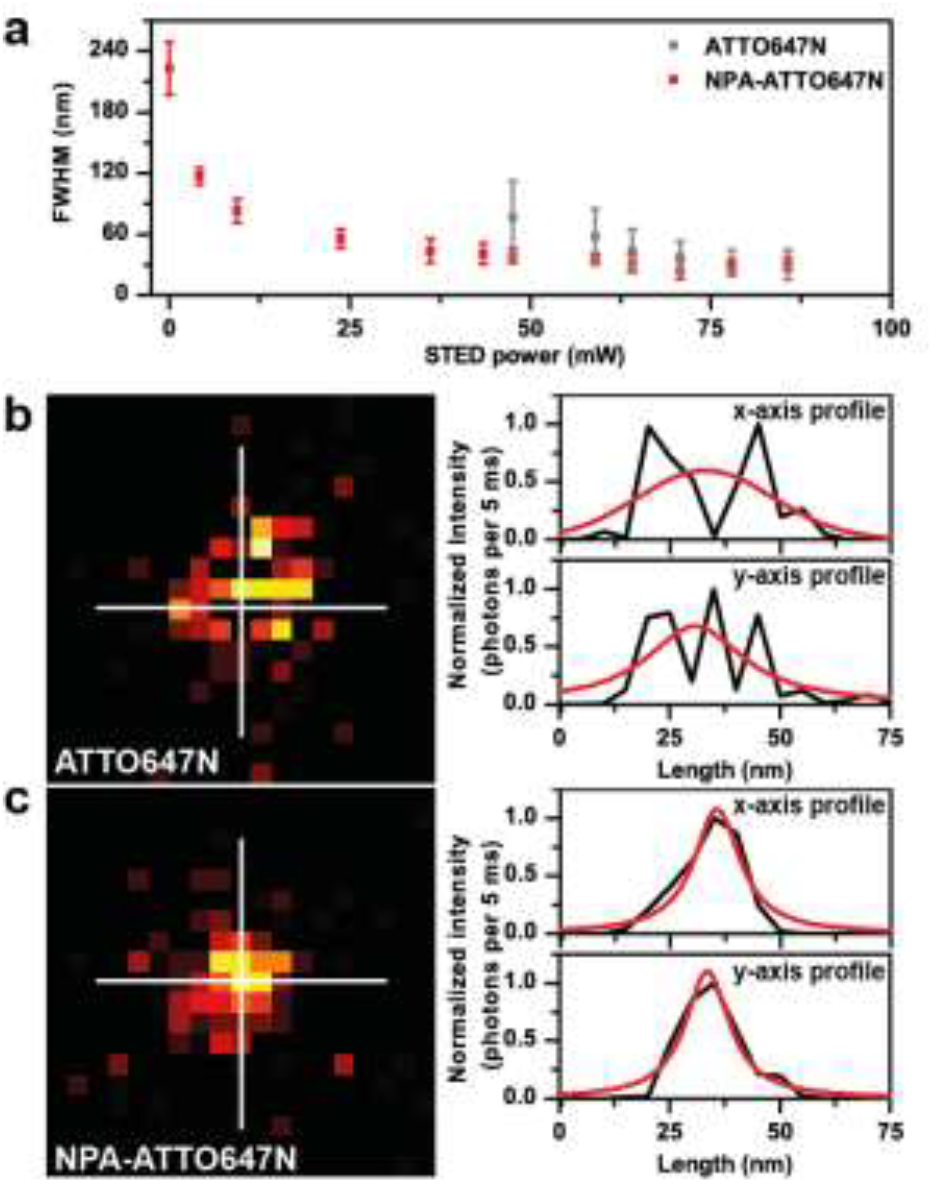
(**a**) Quantitative analysis of the PSF of single ATTO647N conjugates (full width at half maximum, FWHM) at different STED laser powers. The STED images were recorded with 640 nm excitation at 10 kWcm^-2^ complemented with a STED beam at 765 nm with varying powers at the sample. (**b**, **c**) STED images of single ATTO647N (**b**) and NPA-ATTO647N labelled oligonucleotides (640 nm excitation at 3.4 kWcm^-2^ complemented with a STED beam at 765 nm with 50 mW at the sample). All data was recorded with a sample in aqueous PBS buffer at pH 7.4 in the absence of oxygen.

Overlaying the confocal excitation beam with the STED beam results in an increase in the resolution, which is reflected in a decrease in the FWHM of the point-spread-function, PSF (Figure 3a). The optimal resolution was achieved for powers >50 mW and a PSF of 29 ± 5.6 nm (for STED conditions the point spread functions were fitted using a two-dimensional Lorentzian function^37–39^, Figure 3a), which is consistent with values found in literature^10^. For each STED power >20 molecules were analysed using a two-dimensional Lorentzian fit. At high intensities of the STED laser >40 mW, the found FWHM of NPA-ATTO647N was slightly lower compared to ATTO647N (Figure 3a). Additionally, the variation of the FWHM at a specific STED laser intensity was higher for ATTO647N without any photostabilizer. Both observations are likely attributed to the triplet-induced blinking while scanning (Figure 3b). From the images in Figure 3b and c it becomes clear that ATTO647N (Figure 3b) shows multiple pixels with no fluorescence caused raster scanning and blinking in between different positions. This effect is not observed for NPA-ATTO647N, where a single homogenous fluorescent spot was seen (Figure 3c). This observed subtle difference in resolution might, however, have a real world equivalent in STED imaging since the signal from one emitter is not homogenous. The observed saturated resolution decrease > 50 mW can be attributed to limitations of the experimental setup and the length of the dsDNA construct used here as also reported before.^10^

STED microscopy has become a useful tool for high-resolution imaging and our previously published studies with KK114/STAR-RED^24^ and here ATTO647N suggest the potential of photostabilizer-dye conjugates for time-lapse STED-imaging. To probe the available photon-output from individual molecules under STED conditions, something that is potentially useful for dynamic studies such as STED-FCS^40^ of densely-labelled environments, ATTO647N and NPA-ATTO647N were compared in a dynamic imaging mode (Figure 4).

**Figure 4.**
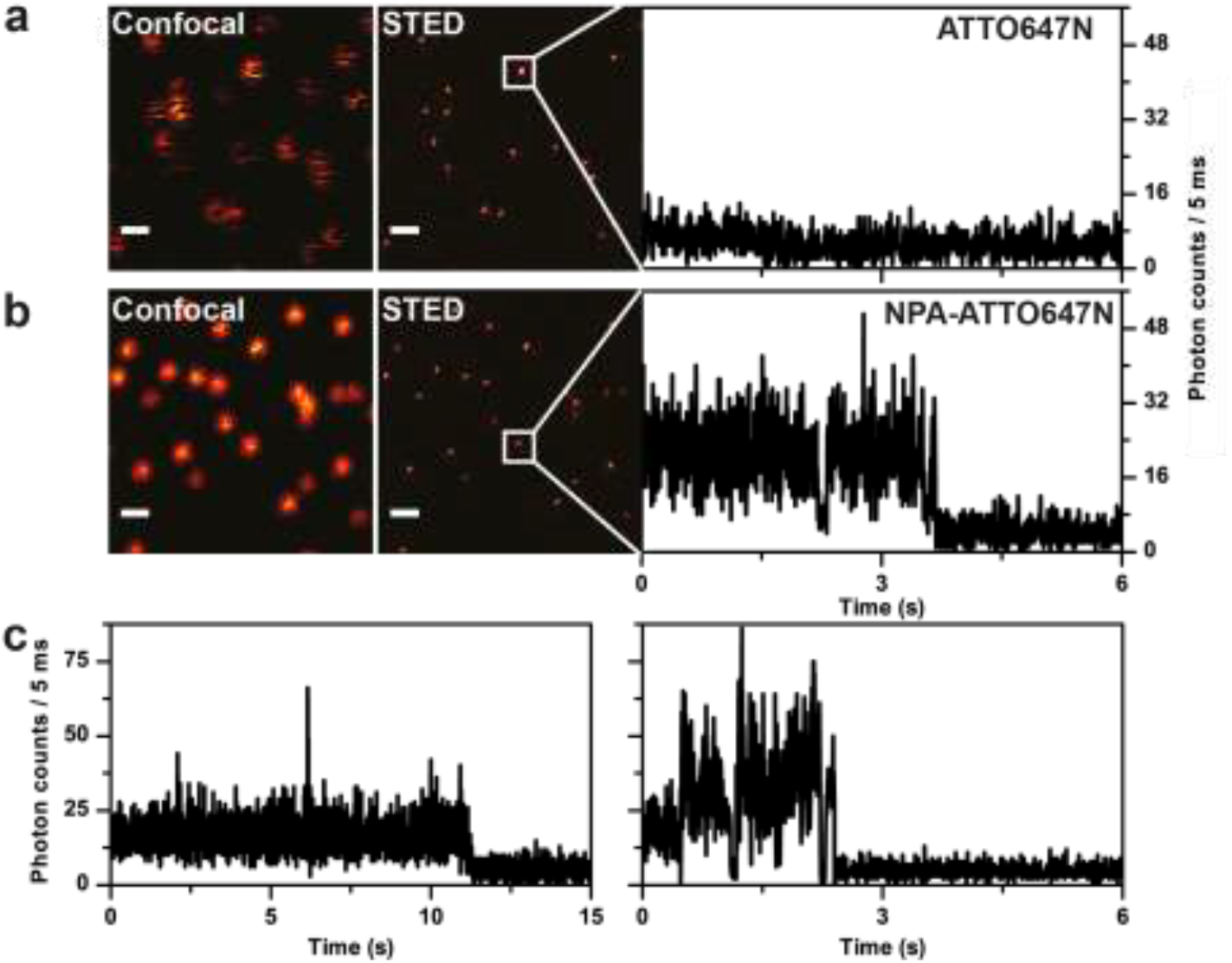
Single molecule fluorescence time traces under STED conditions for (**a**) ATTO647N and (**b**) NPA-ATTO647N. Each of rows consists of a confocal image, STED image and single molecule fluorescence time trace recorded under STED conditions. (**c**) Additional single molecule fluorescence time trace of NPA-ATTO647N recorded under STED conditions. The confocal images were recorded at 640 nm excitation and 3.4 kW cm^-2^. The STED images and single molecule fluorescence traces were recorded with 640 nm excitation at 3.4 kWcm^-2^ complemented with a STED beam at 765 nm with 50 mW at the sample. Scale bars are set to 500 nm. All data was recorded with a sample in aqueous PBS buffer at pH 7.4 in the absence of oxygen.

Next to STED images, a representative single molecule fluorescence time trace is shown under conditions where the STED beam is activated. For ATTO647N no fluorescence signal could be observed under these conditions (Figure 4a). On the contrary, for NPA-ATTO647N fluorescence signals could be observed that persisted for multiple seconds (Figure 4b/c). The results demonstrate that intramolecular photostabilization of organic fluorophores effectively enhances the photophysical properties of fluorophores for STED-type super-resolution microscopy and enables dynamic imaging modes. We believe that the use of self-healing dyes for STED microscopy can be extremely beneficial especially in combination with recent developments like protected STED^41^.

### Photostabilizer-dye conjugates for localization-based STORM microscopy

Thus far we were able to demonstrate that self-healing dyes have an increased photostability compared to their unstabilized counterparts^33^, something that is particularly useful for STED-type microscopy where the number of possible excitation cycles limits the attainable resolution.^32^ For STORM/PALM type super-resolution, however, additional functionality is required. In particular the emission pattern of one fluorophore needs to display photoswitching kinetics with a characteristic low ON/OFF ratio.^42^ This allows to keep most fluorophores inactive and to separate individual labels in a structure via single-molecule localization^1–5,32,43^.

Related to this requirement, our group recently reported the effects of intramolecular photostabilizers on the action of solution-based additives such as Trolox, COT, TCEP and cysteamine (MEA).^44^ In that paper, we evaluated the competition between inter- and intramolecular triplet-state quenching processes and showed that those are neither additive nor synergistic. Our findings revealed that intramolecular processes dominate the photophysical properties of different organic fluorophores for combinations of covalently-linked and solution-based photostabilizers and importantly also photoswitching agents used for STORM. In that study^44^ we identified a new function of intramolecular photostabilizers, i.e., protection of fluorophores from reversible photoswitching. Additionally, evidence was provided that the biochemical environment, here proximity of aromatic amino-acids such as tryptophan, significantly reduced the photostabilization efficiency of commonly used buffer cocktails.^44^

Here, we detail the effects of the intramolecular photostabilizers TX, NPA and COT for Cy5 photoswitching kinetics and photon-yields, i.e., parameters relevant for STORM-type microscopy. We used an established approach where single-molecule TIRF-microscopy provides information on photobleaching lifetime, count-rate, signal-to-noise ratio and total photon-count of individual fluorophores tethered to dsDNA on a microscope coverslide (Figure 5A)^23,44^. It has to be stressed out that the time-point where signal-loss occurs (see Figure 5a, where the signal abruptly decreases to background level) might reflect either irreversible photobleaching or UV-reversible off-switching. In line with our previous work we refer to this as “apparent photobleaching lifetime”^44^, which reflects both pathways for fluorescence deactivation.

**Figure 5.**
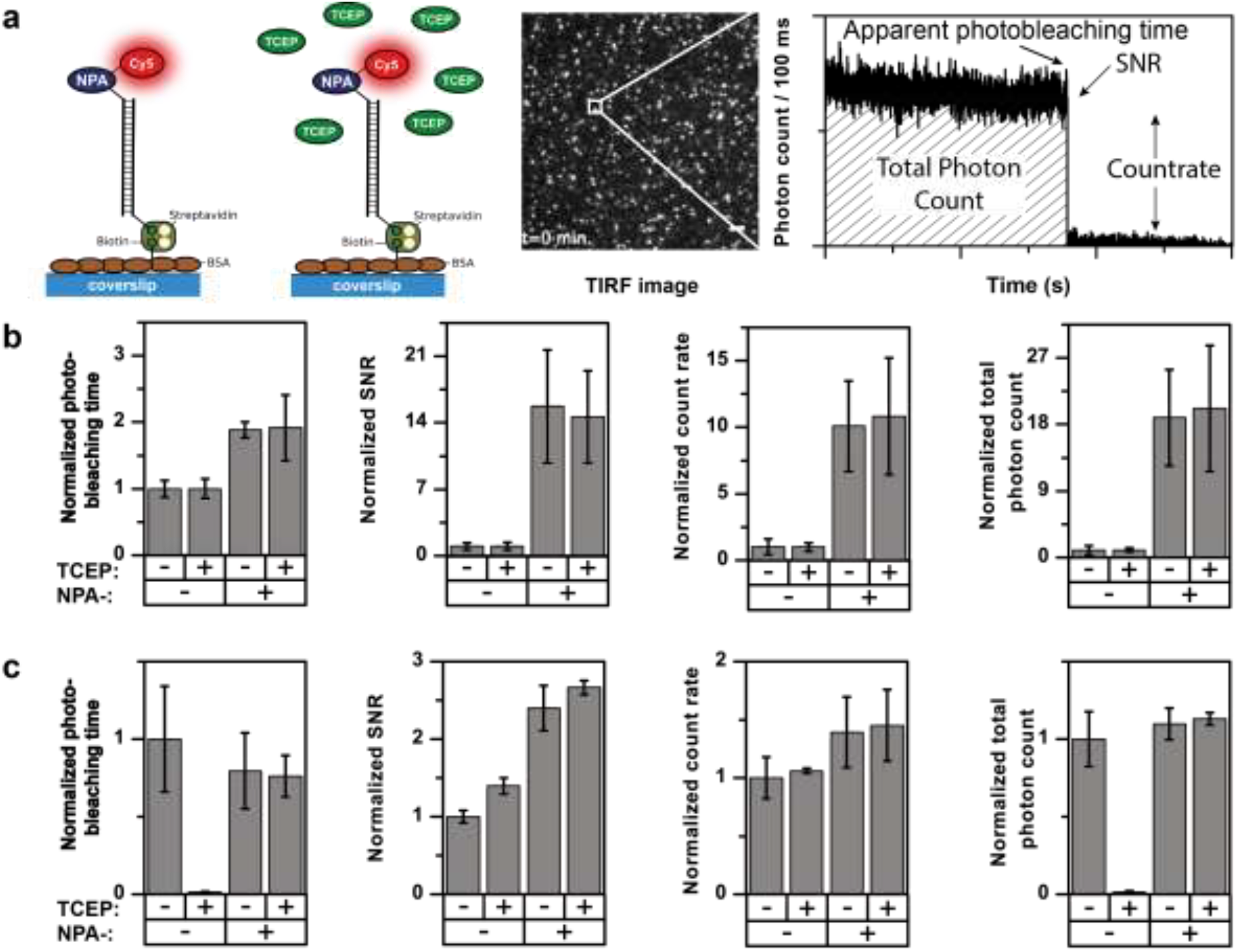
TIRF imaging of Cy5 and ATTO647N derivatives to determine photostability in the presence and absence of TCEP. **(a)** Schematic of the experimental setup of dsDNA-NPA-Dye constructs immobilized via BSA-biotinylated coverslips and a typical fluorescent time-trace obtained in TIRF microscopy. Photophysical characterization of (**b**) ATTO647N and NPA-ATTO647 (**c**) Cy5 and NPA-Cy5 in the absence or presence of 200 μM TCEP. All imaging was done under deoxygenated conditions with 400 W cm^-2^ excitation at 637 nm. Error bars show standard deviation of repeats on 3 different days. Data in the figure partially adapted from Smit *et al.* ^44^.

Since our main interest was the comparison of data in the presence and absence of TCEP in the imaging buffer (Figure 5A), we provide normalized values of all parameters (Figure 5b/c). As shown previously^44^, already lower concentrations TCEP can have a major influence on the photophysical properties of different organic fluorophores (Figure 5b/c). Interestingly, carbopyronines were not influenced in their photophysical properties when TCEP is added to the solution (Figure 5b), which renders them unsuitable for STORM-microscopy in combination with TCEP. As expected from previous work of the Zhuang lab^45^ Cy5 undergoes rapid photoswitching once TCEP is present. Unexpectedly, 200 μM are sufficient to switch-off Cy5 efficiently, while SNR and count-rate are unaltered (Figure 5c).^44^

Upon addition of TCEP, the fluorescence of Cy5 is quenched due to a 1,4-addition of the phosphine to the polymethine bridge of Cy5^46^. It can indeed be seen from Figure 5b,c and Figure 6b,c that the presence of TCEP results in faster signal loss of Cy5 molecules.

**Figure 6:**
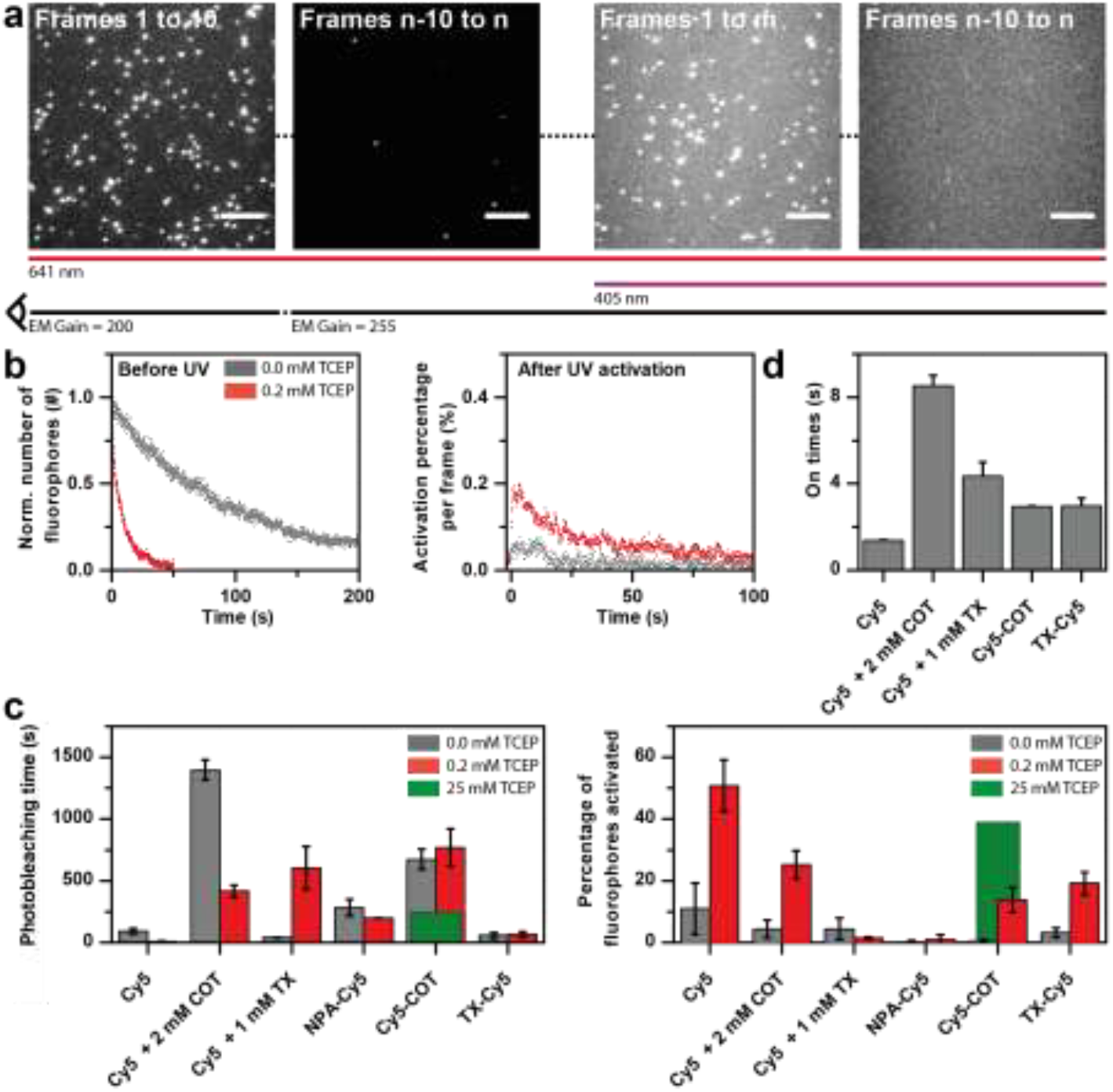
Photoswitching and UV-activation of Cy5 derivatives with TCEP. (**a**) Total Internal Reflection Fluorescence Microscopy images at different time points and under different illumination conditions. (**b**). Decay curve (left) and activation curves (right) of the number of Cy5 fluorophores without (grey) and with (red) TCEP. (**c**) Histograms of the photobleaching time (left) and activation percentages (right) of different cy5 conjugates with (red) and without TCEP (red, green). (**d**) On-times (ms) of the different Cy5 conjugates. The on-times are extracted from the fluorescent time traces. Data in the figure partially adapted from Smit *et al.* ^44^.

In the absence of TCEP the decay is a result of photobleaching whereas in the presence of TCEP the ~10x faster decay is attributed to photoswitching by TCEP (Figure 5c). Subsequently, illumination of UV dissociates the covalent adduct to restore (and activate) 51 ± 8 % of the Cy5 fluorophores (See Figure 6b,c and Vaughan *et al.*^46^), which we also investigated by TIRF microscopy. We determined the percentage of activation using the number of fluorophores in the first frames 1 to 10 at the start of each experiment (to account for blinking molecules, Figure 6a), the remaining number of spots after apparent photobleaching (Frames n-10 to n) and the number of molecules that are activated over the course of illuminating the surface area with UV (405 nm, frames 1 to m), where the number of frames used depends on the photobleaching time of the fluorophore (Figure 6a).

From Figure 6c it is clear that there is little activation of Cy5 in the absence of TCEP. We consequently sought to determine the effect of additional buffer additives COT and TX next to TCEP to minimize blinking and increase the photon yield (Figure 6c)^4^. The addition of COT to Cy5 in the absence of TCEP results in an increased signal duration (Figure 6c, grey bars). The presence of TCEP next to COT quenches Cy5 (with an observed photobleaching lifetime that is ~3x faster) of which 25±5 % could be activated upon illumination with UV. In contrast, the combination of TX and TCEP does not result fast off-switching and high UV-reactivation yield of Cy5. Instead it was observed that photobleaching occurs substantially faster in the absence of TCEP with TX. TCEP thus seems to be required for photostabilization using TX. These results also support the complexity of stabilization mechanisms for Trolox via oxidation^47^ and geminate recombination pathways^48^. We hypothesize that there is a competition between TCEP as an photoswitching agent and as photostabilizer when used in combination with TX. In this specific combination the photostabilization pathway seems more dominant, which is also reflected in the activation percentage showing no significant activation for TCEP and TX (Figure 6c).

In a second step, TCEP quenching was used in combination with photostabilizer-Cy5 derivatives (NPA-Cy5, COT-Cy5 and TX-Cy5). As can be seen in Figure 6c, there is no significant difference between the photobleaching lifetimes in the absence or presence of TCEP for all three constructs. However, the activation percentages do show that activation occurs only in the presence of TCEP. For NPA-Cy5 the activation in the presence of TCEP is negligible but significant for COT-Cy5 and TX-Cy5 with activation of 14 ± 4 % and 19 ± 4 %, respectively.

This apparent contradiction between the unaltered photobleaching times and the activation for COT/TX-Cy5 may be explained by the use of a low concentration of TCEP. At 200 μM TCEP there is a competition between photobleaching of the photostabilizer-Cy5 conjugates and photoswitching *via* TCEP. As we could show before, TCEP has an equilibrium contribution of photoswitching but additionally appears to by switched-off through the triplet state of the fluorophore^44^. In the photostabilizer-Cy5 derivatives however, TCEP-based triplet-quenching (and resulting off-switching) and bleaching are cancelling each other, a fact that results in unaltered photobleaching lifetimes in the presence and absence of TCEP. Consequently, UV-induced on-switching can only be observed with TCEP.

It is therefore hypothesized that when an increased concentration of TCEP was used, the apparent photoswitching pathway *via* TCEP would become more prominent and consequently results in an increased activation percentage upon illumination with UV. To test this hypothesis the concentration of TCEP was increased to 25 mM. In Figure 6c it is shown an increased TCEP concentrations results in reduced photobleaching lifetime of Cy5-COT (i.e., faster off-switching). Upon applying UV-illumination, the number of activated Cy5-COT fluorophores is indeed (Figure 6c) increased compared to 200 μM TCEP. This can be interpreted as that a higher concentration of TCEP results in more photoswitching of Cy5-COT. This could also apply for conditions of Cy5 + 2 mM COT, where COT is efficiently quenching the fluorophores triplet. Looking at the apparent on-times of the different fluorophores upon simultaneous excitation and UV-illumination, it is observed that the on-times increase upon applying photostabilization. Therefore the photon yield per on-period increases. It can be seen that with COT as a photostabilizer in solution the highest on-times are observed. Nevertheless, for intramolecular photostabilized constructs the on-times increase significantly compared to Cy5. As a result the number of photons coming from a single fluorophore, which, as explained previously, is needed to make STORM/PALM techniques useful, increases per on-cycle. The presented trends might allow to establish general trends and to predict better how intramolecular photostabilizers affect photoswitching kinetics for common buffer additives.

Finally, we also tested cysteamine (MEA) as a photoswitching agent for Cy5, in combination with 2 mM COT in solution and self-healing Cy5-COT. Figure 7 shows that upon use of 5 mM MEA the apparent photobleaching lifetime is reduced by MEA compared to the use of TCEP. This is also true when used for Cy5 in combination with COT as a photostabilizing agent in the imaging buffer. However, looking at the activation percentages upon applying UV, it can be seen that only for Cy5 in the absence of COT, also the activation percentage has increased further when using MEA. For all other conditions the activation efficiency of MEA is lower compared to TCEP.

**Figure 7:**
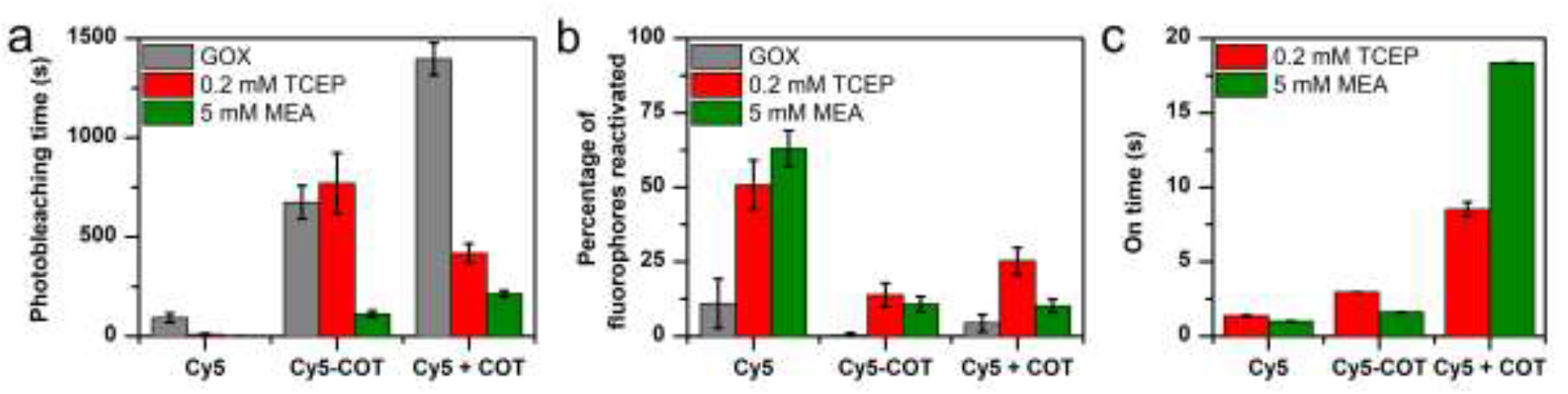
Comparison of Cy5 with inter- and intramolecular photostabilized with COT with photodarkening by 0.2 mM TCEP and 5 mM MEA. (**a**) Photobleaching lifetime (**b**) Percentage of fluorophore recovered by 405 nm excitation. (**c**) Mean on-state lifetime of reactivated fluorescent transients in case of reactivation. Error bars are standard deviation of independent repeats of different days (**a**, **b**) or SEM (**c**). Data in the figure partially adapted from Smit *et al.* ^44^

Finally, we studied the on-times after photoactivation for molecules/conditions with reactivation efficiency> 15%. Interestingly, the on-time for the condition of Cy5 with COT in solution are doubled when cysteamine is present, thereby increasing the photostability of the fluorophore. The on-times for Cy5 and Cy5-COT were reduced when cysteamine is added. Consequently, the photon-yield coming from a single molecule upon simultaneously applying excitation and UV-illumination is decreased making it less useful for STORM/PALM techniques.

## 4. Discussion and conclusions

Our results show that intramolecular photostabilization of organic fluorophores effectively enhances the photophysical properties of fluorophores for STED microscopy. Photostabilizer-dye conjugates show an increased photostability and brightness, which are important parameters for *in vivo* and *in vitro* STED microscopy applications. Upon covalent binding of a nitrophenyl moiety to ATTO647N, single molecule count rates of up to MHz were obtained. It was found that intramolecular photostabilization significantly improves the photostability of ATTO647N when excited with both the confocal excitation and STED beam, thereby increasing the number of possible subsequent STED images that can be acquired. Additionally, on average lower STED laser intensities were needed to obtain the maximum resolution ~35 nm with NPA-ATTO647N when compared to ATTO647N, thereby reducing the probability of photobleaching of the fluorophores and phototoxicity to the sample. The improvement of the photophysical parameters via intramolecular photostabilization paves the way for dynamic STED imaging without the need of adding (potential toxic) chemical compounds.^10^

In the second part of this contribution it was shown how intramolecular photostabilizers affect the photoswitching kinetics of TCEP and MEA in combination with Cy5. It was shown that both TCEP and MEA cause reversible Cy5 off-switching, allowing reactivation upon UV illumination. However, by adjusting the TCEP concentration the reactivation percentage and on-times of photostabilized conjugates could be optimized allowing their application in STORM-type super-resolution.

When using MEA as a photoswitching agent, we find a reduced apparent photobleaching lifetime which is not translated into a corresponding increase in reactivation percentage. The reduction in photobleaching time can therefore not be attributed to the formation of a MEA-induced reversible dark state. This discrepancy is only observed when COT is present in both inter- and intramolecular photostabilization, which suggests that either MEA interferes with reactive intermediates in the mechanism of photostabilization, or in the presence of photostabilization a different dark state is formed which is less susceptible to UV-induced photoactivation.

The longest on-times are observed when using MEA with COT as a photostabilizer in solution, which suggests that under these conditions there is the highest potential of collecting the most photons per activation. Therefore, further investigation should be aimed towards identifying potential interactions between COT photostabilization and thiol-based photoswitching agents, which can provide guidance in optimizing the reactivation efficiency when COT is employed as a photostabilizer.

## 5. Material and methods

### 5.1. Sample preparation and surface immobilization of oligonucleotides

Immobilization and the study of single fluorophores was achieved using a dsDNA scaffold comprising two 40-mer oligonucleotides, i.e. ssDNA-fluorophore and ssDNA-biotin. Sequences of both oligomers were adapted from literature^47,49,50^. Single immobilized fluorophore molecules were studied in LabTek 8-well chambered cover slides (Nunc/VWR, The Netherlands) with a volume of 750 μl, as described in literature^51^. After cleaning with 0.1% HF and washing with PBS buffer (one PBS tablet was dissolved in deionized water containing 10 mM phosphate buffer, 2.7 mM potassium chloride and 137 mM sodium chloride at pH 7.4; Sigma-Aldrich, The Netherlands), each chamber was incubated with a mixture of 5 mg/800 μL BSA and 1 mg/800 μL BSA/biotin (Sigma Aldrich, The Netherlands) at 4°C in PBS buffer overnight. After rinsing with PBS buffer, a 0.2 mg·ml^-1^ solution of streptavidin was incubated for 10 minutes and subsequently rinsed with PBS buffer.

The immobilization of dsDNA was achieved via a biotin-streptavidin interaction using pre-annealed dsDNA with the aim of observing single emitters for prolonged periods of time and allowing free rotation of the fluorophores. For this, 5-50 μL of a 1 mM solution of ssDNA-fluorophore or ssDNA-NPA-fluorophore (ATTO647N-(NPA)-C6-5’-TAA TAT TCG ATT CCT TAC ACT TAT ATT GCA TAG CTA TAC G-3’, used as received from Eurofins and IBA, Germany) was mixed with the complementary ssDNA-biotin at the same concentration (biotin-5’-CGT ATA GCT ATG CAA TAT AAG TGT AAG GAA TCG AAT ATT A-3’, used as received from IBA, Germany). The respective mixtures of the two oligomers were heated to 98°C for 4 minutes and cooled down to 4°C with a rate of 1°C·min-1 in annealing buffer (500 mM sodium chloride, 20 μM Tris-HCL, and 1 mM EDTA at pH = 8). The treated LabTek cover slides were incubated with a 50-100 pM solution of pre-annealed dsDNA for 1-2 minutes. All single molecule experiments were carried out at room temperature (22 ± 1°C). Oxygen was removed from the buffer system utilizing an oxygen-scavenging system (PBS, pH = 7.4, containing 10% (wt/vol) glucose and 10% (vol/vol) glycerine, 50 mg· ml^-1^ glucose-oxidase, 100-200 μ g· ml^-1^ catalase. Glucose-oxidase catalase (GOC)^51,52^ was used instead of a combination of protocatechuic acid and protocatechuate-3,4-dioxygenase (PCA/PCD)^53^ to avoid convolution of inter- and intramolecular photostabilization with PCA^54^.

The procedures for the functionalization of oligonucleotides have been established before and described in detail in Smit *et al.* ^44^.

### 5.2 Confocal scanning microscopy and data analysis

A custom-built confocal microscope, described in literature^23^, was used to study the fluorescence properties of organic fluorophores on the level of single molecules. Excitation was achieved via a spectrally filtered laser beam of a pulsed supercontinuum source (SuperK Extreme, NKT Photonics, Denmark) with an acoustooptical tunable filter (AOTFnc-VIS, EQ Photonics, Germany), leading to 2 nm broad excitation pulses centred around 640 nm. The spatially filtered beam was coupled into an oil-immersion objective (60×, numerical aperture (NA) 1.35, UPLSAPO 60XO mounted on an IX71 microscope body, both from Olympus, Germany) by a dichroic beam splitter (zt532/642rpc, AHF Analysentechnik, Tuebingen, Germany). Surface scanning was performed using a XYZ-piezo stage with 100 × 100 × 20 μm range (P-517-3CD with E-725.3CDA, Physik Instrumente, Germany). Fluorescence was collected by the same objective, focused onto a 50 μm pinhole and detected an avalanche photodiode (t-spad, >50 dark counts per second, Picoquant, Germany) with appropriate spectral filtering (ET700/75 AHF, both from Analysentechnik). The detector signal was registered using a Hydra Harp 400 ps event timer and a module for time-correlated single-photon counting (both from Picoquant). The data was evaluated using custom-made LabVIEW software^49,50^ Blinking kinetics were extracted from fluorescent time traces in the form of ON and OFF times according to established procedures^49^. Fluorescence lifetimes were determined using time-correlated single-photon counting as described before^23^.

### 5.3 Single molecule STED and confocal microscopy

Single molecule STED microscopy and the corresponding confocal microscopy were performed on a Microtime-200 STED microscope (PicoQuant, Berlin, Germany). Excitation was performed at 640 nm (LDH-D-C-640P laser diode, PicoQuant) and STED at 765 nm, at 10 MHz. A donut-shaped intensity distribution of the STED laser was realized by the easy-STED approach, that is, a /4 phase plate was inserted into the collimated STED beam, which is overlaid to the excitation beam using the appropriate filter (SBDC 565, AHF, Analysentechnik, Germany). Both the excitation and STED laser were focussed onto the sample via an oil-immersion objective (UPlanSapo 100×, NA 1.4, Olympus, Japan) mounted on an IX73 microscope body (Olympus, Germany) by dichroic beam splitter (ZT 640/752 rpc-UF3, Chroma, USA). Fluorescence was collected through the same objective and detected by avalanche photodiodes (Excelitas Technologies, Quebec, Canada) with corresponding spectral filtering (HQ 690/70, AHF Analysentechnik, Germany). The data was analysed with Symphotime64 software (PicoQuant, Berlin, Germany). For fitting the point spread functions with a 2-dimensional Lorentzian and/or Gaussian function a custom made ImageJ macro was used. The following 2-dimensional Gaussian function was used: 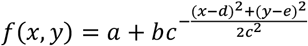, and the following 2-dimensional Lorentzian function was used: 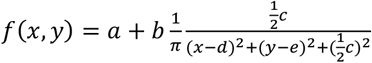 with *a* as background, *b* the aplitude, *c* the spread (sigma), *d* the x position and *e* the center y position.

### 5.4. STORM microscopy

Widefield TIRF imaging was performed on an inverted microscope (Olympus IX-71, UPlanSApo x100 NA 1.49 Objective, Olympus, Germany) in a similar manner as described before^23^. Images were collected with a back-illuminated electron multiplying charge-coupled device camera (512×512 pixel, C9100-13, Hammamatsu, Japan) with matching filters and optics. To study the reactivation and STORM parameters, the sample was illuminated with a 375nm or 405nm laser after photodarkening.

Individual fluorophores were detected in TIRF movies using a fixed threshold and discoidal averaging filter. The number of emitters as a function of time was fitted to a mono-exponential decay to obtain the mean photobleaching or photodarkening lifetime. To determine the reactivated fluorophores again a fixed threshold and discoidal averaging filter was used over the first n frames after the start of illuminating with both the excitation and UV light. For a typical experiment, 5 movies were recorded of a given condition, which was repeated on 3 different days. Fluorescent transients were extracted from the data by selecting a 3×3 pixel area (pixel size 160 nm) around the emitter and plotting the resulting mean fluorescence intensity in time. These fluorescent transients were then processed in home-written software to extract other photophysical parameters such as signal-to-noise ratio and count-rate.

## Acknowledgements

This work was financed by an ERC Starting Grant (ERC-STG 638536 – SM-IMPORT to T.C.). The Zernike Institute for Advanced Materials and the Centre for Synthetic Biology (University of Groningen) provided funding for purchasing a STED microscope. J.H.M.v.d.V acknowledges the Ubbo-Emmius funds (University of Groningen) for a PhD. stipend. T.C. was supported by the Center of Nanoscience Munich (CeNS), Deutsche Forschungsgemeinschaft within GRK2062, LMUexcellent and the Center for integrated protein science Munich (CiPSM).

